# Testes of *DAZL* null sheep lack spermatogonia and maintain normal somatic cells

**DOI:** 10.1101/848036

**Authors:** Zachariah McLean, Sarah Jane Appleby, Jingwei Wei, Russell Grant Snell, Björn Oback

**Affiliations:** Reproduction, AgResearch, Ruakura Research Centre, Hamilton, New Zealand; Applied Translational Research Group and Centre for Brain Research, School of Biological Sciences, University of Auckland, Auckland, New Zealand; Department of Molecular Medicine & Pathology, School of Medical Sciences, University of Auckland, Auckland, New Zealand

## Abstract

Multiplying the germline would increase the number of offspring that can be produced from selected animals, accelerating genetic improvement for livestock breeding. This could be achieved by producing multiple chimaeric animals, each carrying a mix of donor and host germ cells in their gonads. However, such chimaeric germlines would produce offspring from both donor and host genotypes, limiting the rate of genetic improvement. To resolve this problem and produce chimaeras with absolute donor germline transmission, we have disrupted the RNA-binding protein *DAZL* and generated germ cell-deficient host animals. Using Cas9 mediated homology-directed repair (HDR), we introduced a *DAZL* loss-of-function mutation in male ovine fetal fibroblasts. Following manual single-cell isolation, 4/48 (8.3%) of donor cell strains were homozygously HDR-edited. Sequence-validated strains were used as nuclear donors for somatic cell cloning to generate three lambs, which died at birth. All *DAZL*-null male neonatal sheep lacked germ cells. Somatic cells within their testes were morphologically intact and expressed normal levels of somatic cell-specific marker genes, indicating that the germ cell niche remained intact. This extends the *DAZL-*mutant phenotype beyond mice into agriculturally relevant ruminants, providing a pathway for using absolute transmitters in rapid livestock improvement.

## Introduction

Control of the germline in livestock species by assisted reproductive technologies is valuable for animal breeding by driving genetic gain. Simulations have predicted a gain of several years’ worth of genetic improvement by multiplying the germline of an elite sire, and therefore the number of its offspring, when compared to conventional breeding approaches ^1^. Multiplication can be achieved by inserting the germline of an elite donor into the gonads of several germ cell-deficient hosts, producing chimaeric sires with cells from distinct zygotic origins. To create such sterile hosts, local testes irradiation has successfully depleted germ cells in sheep ^2^ and cattle ^3^. However, this method requires general anaesthesia and access to expensive specialist equipment ^4^. Depending on the radiation dose and animal’s age at time of radiation, it may also leave some endogenous germ cells intact ^2^, compromise testicular somatic cells ^5–7^, slow down recovery of spermatogenesis and even reduce libido ^7^. The alternative strategy of sterilisation by fetal busulfan treatment resulted in systemic toxicity in pigs and germ cell ablation was again incomplete ^5^. Resulting sperm output would be a mixture of donor and host cells, which is unacceptable for breeding applications that require absolute transmission of the elite donor germline. Therefore, production of a suitable host animal by physical or chemical means remains sub-optimal.

Recently, disruption of key germ cell development genes *NANOS2* and *NANOS3* has ablated germ cells in pigs and cattle, respectively ^8,9^. *NANOS3* was disrupted in female cattle, but *NANOS2* was knocked out in both sexes, with only male germ cells being affected. Both knockouts reproduce the mouse phenotype and provide a suitable genetically sterilised host animal completely devoid of endogenous germ cells ^10^. As an alternative target, the RNA-binding protein Deleted in Azoospermia-(DAZ)-Like (*DAZL*) is highly conserved among vertebrates, including bony fish, chicken, mice and humans ^11^. DAZL functions by post-transcriptionally binding mRNA in 3′ untranslated regions, acting either as a transcriptional repressor or activator in a network regulating the cell cycle and the maintenance of germ cell identity ^12^. In mice lacking *Dazl*, primordial germ cells (PGCs) fail to commit to a germ cell lineage and maintain their undifferentiated state ^13^. This leads the aberrant germ cells down an apoptotic pathway, producing adult mice of both sexes that entirely lack germ cells ^14,15^. In livestock, *DAZL* has been targeted in pigs, confirming complete ablation of the germline.^16,17^.

Although introducing a loss-of-function modification by classical gene targeting is possible in livestock, such as the approach described for producing *NANOS3* mutant cattle, the efficiencies are low ^18^. Programmable nucleases, like TALENs or Cas9, can introduce precise double strand breaks at desired loci and greatly enhance the efficiency of gene targeting. This was demonstrated in pigs with TALENs targeting *DAZL* and Cas9 targeting *NANOS2* ^8,16^. To generate animals, introducing genome editors at the zygote stage is a viable approach, but this typically leads to complex genetic mosaics with an unpredictable and difficult-to-define phenotype in the first generation ^19,20^. In mosaic animals, the germline would be formed by an unknown proportion of functional wild-type cells, complicating retrospective determination of the extent of germ cell loss due to lack of *DAZL*. By contrast, animals produced by somatic cell cloning will be non-mosaic as they derive from a single genome-edited donor nucleus. Therefore, somatic cell cloning enables accurate phenotyping of *DAZL*-null offspring in the first generation. This allows a strategy for chimaerising *DAZL*-null host with wild-type donor cloned embryos, generating chimaeric animals which transmit only the donor germline (“absolute transmitters”).

Here we used gRNA-Cas9 mediated homology-directed repair (HDR) to produce a *DAZL* loss-of-function mutation in male ovine fetal fibroblasts. Following manual isolation of edited donor cell strains and somatic cell cloning, we generated *DAZL*-null male neonate sheep devoid of germ cells. This extends the *DAZL-*mutant phenotype beyond mice and pigs, providing a pathway for the use of absolute transmitters in accelerated livestock improvement.

## Results

### gRNA-Cas9-mediated disruption of *DAZL*

To validate the genomic sequence predictions of the ovine *DAZL* gene available from NCBI, we compared the mRNA and protein sequences to the murine and bovine orthologues. Aligning the sequences, the exon structure was similar, but the annotated sheep translation start site differed considerably compared to mice or cattle (Supplementary Fig. S1). For both mice and cattle, there was an ATG translation initiation codon at the 3’ end of exon 1, whereas for sheep, the entire exon 1 of the sheep gene was annotated to be translated. Overall, there was high identity between all three species from exon 2-10, but the 5’ and 3’ untranslated regions were less certain in sheep.

Cas9 was used to introduce a loss-of-function mutation in exon 2, truncating the protein upstream of the predicted RNA recognition motif and DAZ functional domains (Fig. 1a). An HDR template was designed to insert a 6 bp sequence into exon 2 of the *DAZL* gene, comprising a premature stop codon followed by a *TaqI* restriction site for easy detection (Fig 1b). Three potential single guide RNAs (gRNA) were selected with minimal off-target sites using the software package CCTop ^21^. After transfecting ovine fetal fibroblasts (OFFs) with the gRNA-Cas9 expression plasmid and the single-stranded oligonucleotide repair template, non-transfected cells were removed by transient puromycin selection. Quantification by droplet digital (dd) PCR showed that the average number of HDR events across three transfections was higher for gRNA1 with 6.3 ± 3.1%, compared to 0.26 ± 0.13% for gRNA2, and 0.058 ± 0.058% for gRNA3, even though this was not significant when modelled with an ANOVA (*P* = 0.11). A mixed cell population from gRNA1 with 12.6% HDR was subcloned by manual selection of mitotic cells to obtain a pure donor strain for subsequent somatic cloning. Of 118 mitotic cells seeded, 48 (41%) proliferated sufficiently for analysis. A PCR-based allelic discrimination assay on expanded strains showed 5/48 (10%) were positive for only the mutant allele (Supplementary Fig. S2). All putative mutants (5/5=100%) were confirmed as biallelically edited by *Taq*I restriction digest (Supplementary Fig. S3). One of these clones (#43) contained the correct 6 bp insertion on one allele, as well as an approximate 200 bp anomalous insertion around the Cas9 cut site on the other allele. The remaining four strains were validated by Sanger sequencing (Supplementary Fig. S4). Overall, specific HDR-mediated disruption of both *DAZL* alleles was achieved with 4/48 (8%) efficiency.

**Figure 1.**
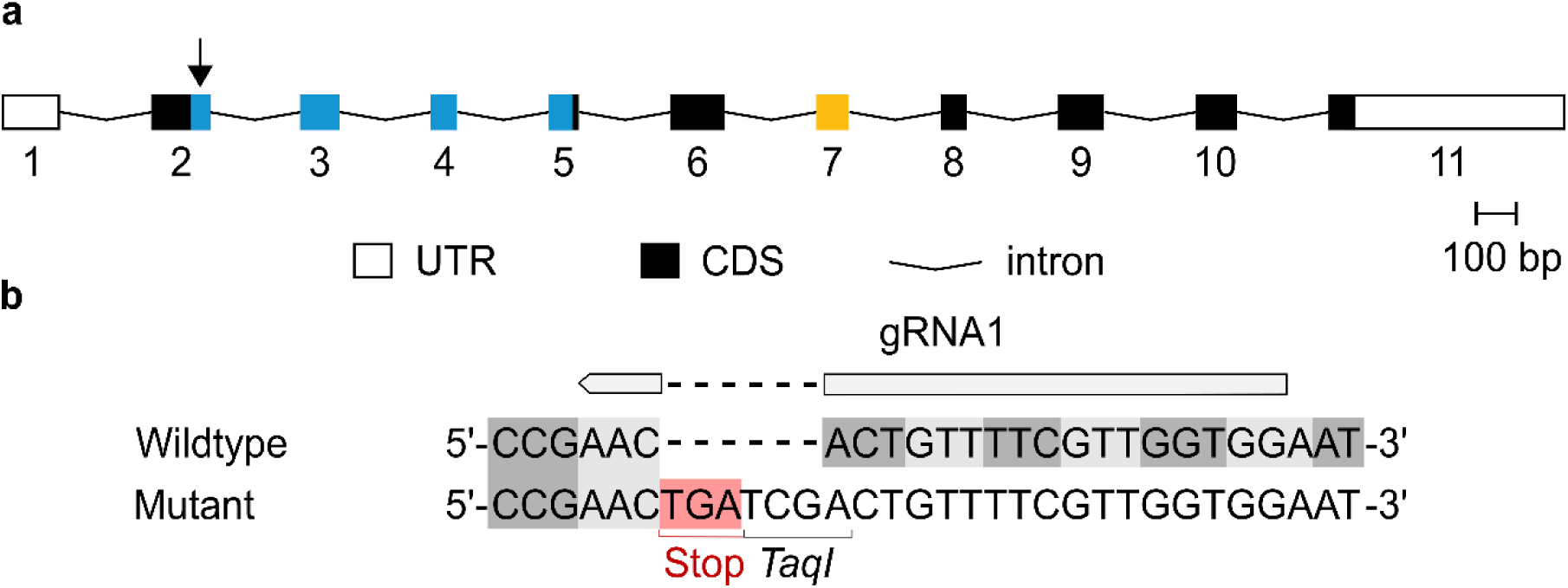
gRNA-Cas9 genome editing of ovine *DAZL*. (a) The ovine *DAZL* gene structure, including numbered exons displaying the predicted untranslated region (UTR) and coding sequence (CDS), with the Cas9 target site indicated by the arrow above exon 2. The RNA recognition motif (blue) and DAZ (yellow) domains were mapped from the mouse sequence. Introns and UTR are not to scale; (b) Cas9 cleavage mediates an HDR insertion of a stop codon and *Taq*I restriction site. Alternating codons are indicated with grey shading to highlight the insertion of the 6 bp mutation in-frame, with the stop codon highlighted in red.

To test specificity of the gRNA-Cas9 system, a biased Sanger sequencing screen of the top three potential off-target sites, as identified by CCTop, revealed no mutations (Supplementary Fig. S5). However, a PCR-based screen for the Cas9 coding region revealed that all four correctly edited clonal strains had integrated the gRNA-Cas9 plasmid into their genome (Supplementary Fig. S6). Chromosome numbers of the four correctly edited OFF3 strains neither differed significantly from low-passage parental wild-type cells, nor from ovine adult fibroblasts (OAF1) that had previously produced viable cloned lambs in our hands (Supplementary Fig. S7) ^22^.

### Generation of cloned *DAZL*-deficient lambs

Homozygous clonal knockout (KO) strains 9 and 25 were randomly chosen as donor cell strains for zona-free somatic cell cloning. Strains 9 and 25 developed into blastocysts at a rate of 17/77 (22%) and 16/90 (18%), respectively, which was comparable to non-edited parental OFFs at 7/35 (20%) (Table 1). Cloned blastocysts on day 7 were transferred into hormonally programmed ewes by laparoscopic surgery, either fresh or after warming from vitrification, with a total of 2/13 (15%) embryos surviving to term for strain 9, and 1/13 (8%) for strain 25 (Table 2). Both ewes carrying lambs from strain 9 developed symptoms of hydroallantois at term. They were induced to deliver at gestational day 147 but did not respond and were therefore euthanised to surgically recover the lambs at day 151. Both recovered lambs died within minutes due to respiratory failure. The lamb from strain 25 was delivered dead on day 146, displaying oedematous and decaying tissues. All lambs that failed at the perinatal stage displayed a variety of anatomical malformations typically observed in somatic cell clones ^23,24^, including ankylosis of the forelimbs and hydronephrosis. Developmental rates of cloned concepti *in vitro* and *in vivo* were well within the range reported for sheep ^25–28^.

**Table 1.** Somatic cell cloning in vitro development of DAZL^-/-^ cloned embryos.

| Donor cells | n<sup>1</sup> | nIVC<sup>2</sup> | ≥ 2-cell<br>(%)<sup>3</sup> | Total<br>blastocysts | Grade 1-2<br>blastocysts |
| --- | --- | --- | --- | --- | --- |
|  |  |  |  | (%)<sup>4</sup> | (%)<sup>5</sup> |
| OFF *DAZL*<sup>+/+</sup> parental line | 1 | 40 | 35 (88) | 7 (20) | 2 (29) |
| OFF *DAZL*<sup>-/-</sup> strain 9 | 2 | 78 | 77 (99) | 17 (22) | 5 (29) |
| OFF *DAZL*<sup>-/-</sup> strain 25 | 2 | 96 | 90 (94) | 16 (18) | 8 (50) |

<sup>1</sup> Number of somatic cell cloning runs.
<sup>2</sup> Number of embryos placed into *in vitro* culture (IVC).
<sup>3</sup> Cleavage normalised on nIVC.
<sup>4</sup> Blastocyst development normalised on ≥ 2-cell embryos.
<sup>5</sup> Proportion of grade 1-2 blastocysts normalised on total number of blastocysts.

**Table 2.**
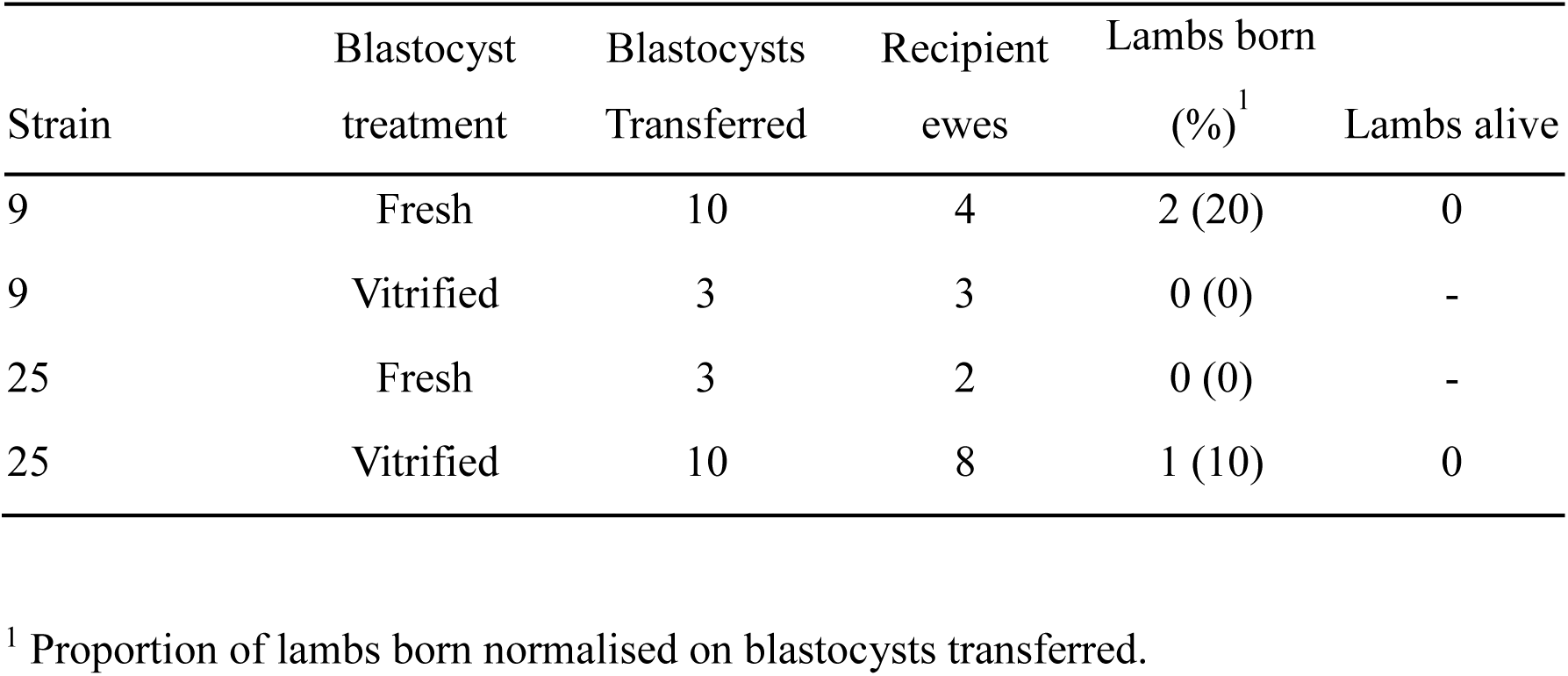
In vivo survival of DAZL^-/-^ cloned embryos.

### *DAZL* mutant phenotype

Testes harvested from the two deceased lambs derived from cell strain 9 were histologically analysed. Compared to *DAZL*^+/+^ controls, *DAZL^-/-^* testis cords comprised somatic support cells but lacked spermatogonia (Fig. 2a). Immunostaining on cryosections revealed a cytoplasmic signal for spermatogonial marker DDX4 in the centre of testis cords for *DAZL*^+/+^ but not for *DAZL^-/-^* lambs (Fig. 2b). These observations were confirmed for different wild-type and KO animals across various locations in the testis (Supplementary Fig. S8).

**Figure 2.**
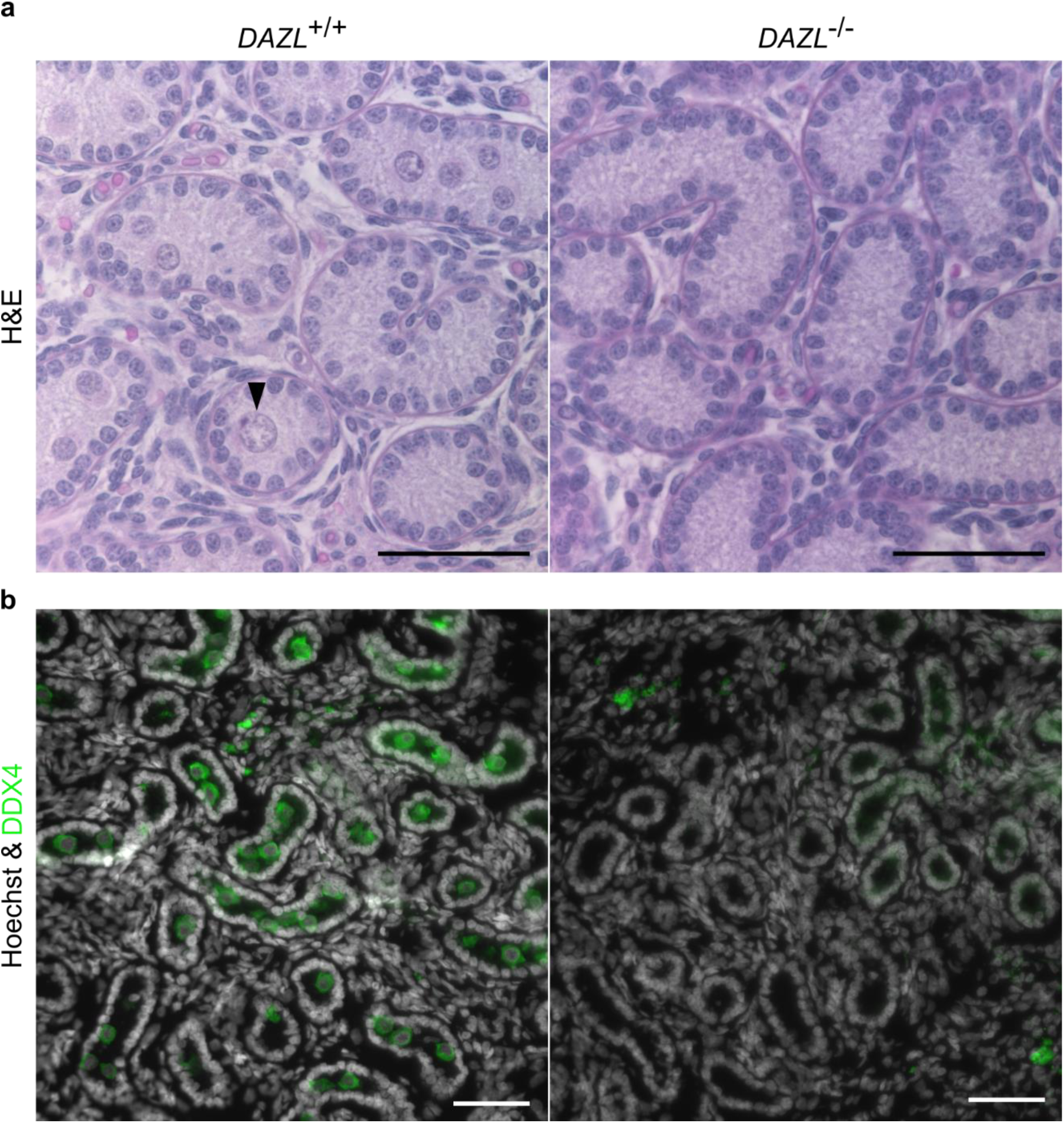
Germ cell-deficient phenotype in *DAZL*^-/-^ neonate testis. (a) Haematoxylin and eosin stained sections of neonate lamb testis. For *DAZL^+/+^,* an arrow indicates a putative spermatogonium in the lumen of a testis cord. (b) Immunostaining of DDX4 in neonate lamb cryosections, co-stained for DNA (Hoechst 33342). Scale bars are 50 µm.

Lineage-specific gene expression was analysed to quantify germ and somatic cell populations in lamb testes by qPCR. Mutant testis parenchyma showed abolished or significantly reduced germ cell-specific expression of *DAZL*, *NANOS2*, *DDX4*, *SALL4*, *LIN28*, and *NANOG* compared to wild type (Fig. 3a). By contrast, discriminatory markers for Sertoli and peritubular myoid cells (*GDNF*), Leydig cells (*CSF1*, *HSD3B1*) and both Leydig and Sertoli cells (*GATA4*, *NR5a1*) were unaffected (Fig. 3b). We conclude that the introduced *DAZL* loss-of-function mutation led to a germ cell-deficient phenotype without compromising development of somatic support cells.

**Figure 3.**
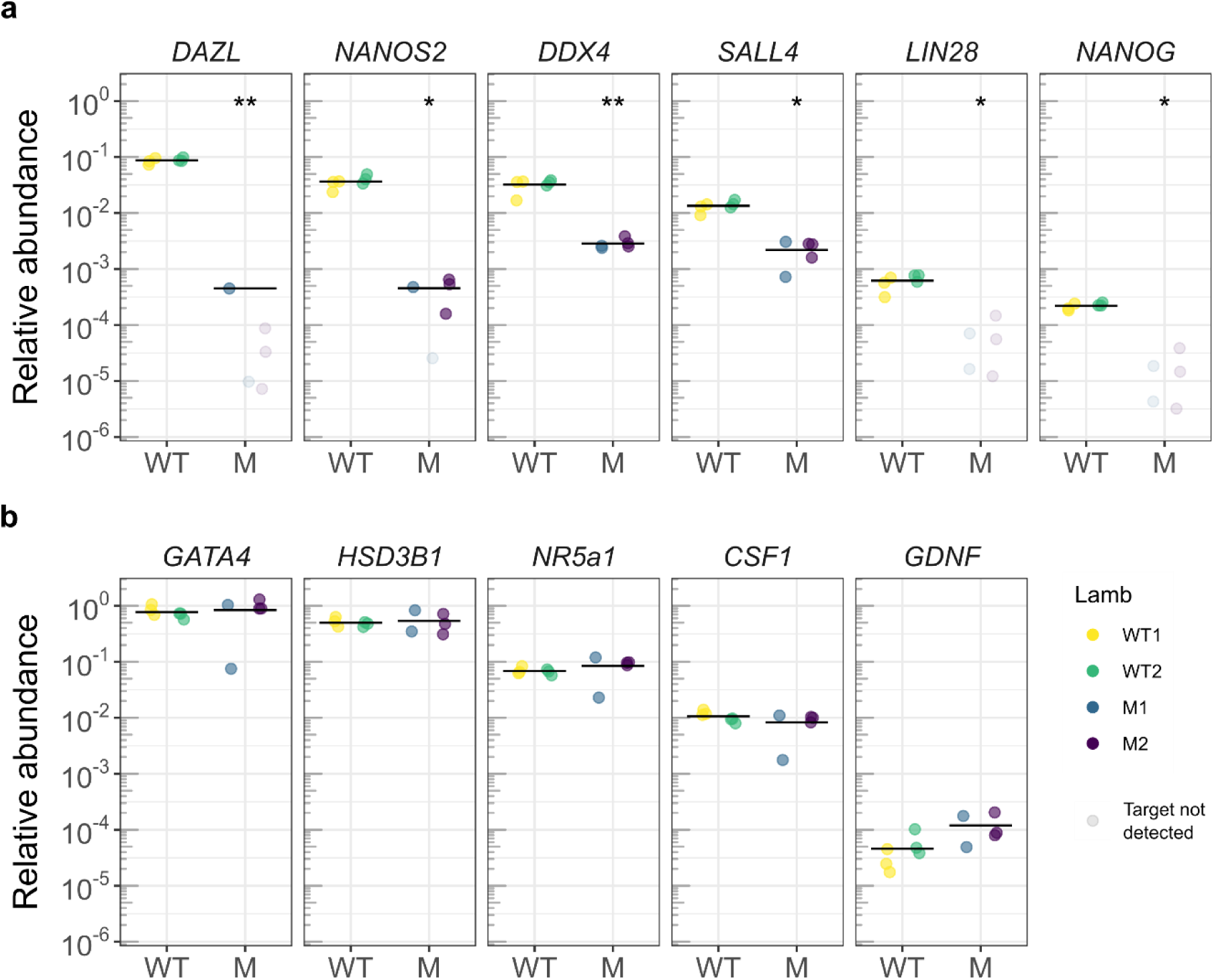
Reduction of germ cell signature but maintenance of somatic expression in *DAZL*^-/-^ testis. Gene expression in testis parenchyma of wild-type (WT) and mutant (M) lambs, showing relative expression of (a) germ cell and (b) somatic markers. Significance values after linear mixed-effects model represented by * *P* < 0.05 and ** *P* < 0.01.

## Discussion

Our data provide another proof-of-principle for using HDR to precisely modify the sheep genome ^29^. This has been shown to be useful for producing human disease models in areas where sheep more physiologically relevant ^29^. The HDR approach was convenient as the appearance of a known allele simplifies the screening of strains. It allows direct insertion of a premature termination codon, which prevents the formation of novel protein sequences with unknown function after NEHJ ^30^. However, caution is required when selecting strains to ensure the expression plasmid has not integrated into the genome, which has been shown to happen with the backbone of HDR constructs for introducing the hornlessness (polled) trait in cattle ^31,32^. Plasmid integration may have been favoured by the transient puromycin selection we applied to enhance editing efficiency. These issues can be avoided by screening lines before somatic cell cloning, or by transfecting CRSIPR-Cas9 as a ribonucleoprotein complex.

Even though low efficiency limits the large-scale production of cloned farm animal models, somatic cell-mediated genome editing provides an effective tool for generating defined genotypes. This is in contrast to embryo-mediated editing that often requires time-consuming breeding strategies to obtain non-mosaic offspring from mosaic founder animals ^19^. Developmental malformations, a known side effect of the cloning procedure, are a potential confounding variable when analysing the *DAZL^-/-^* phenotype. However, the developmental defects associated with somatic clones have not been reported to include germ cells ^23^. This provides confidence in the specificity of the *DAZL^-/-^* phenotype to the germline in sheep. Similarly, the death of the three *DAZL^-/-^* lambs perinatally can be attributed to somatic cell cloning rather than genome editing. This poor viability at the neonatal stage has been observed in most published reports involving sheep clones, with at least half of the term animals perishing within the first day ^26,28^. The main developmental problems in sheep clones have affected the pulmonary, nephrotic, and musculoskeletal systems ^24^. These abnormalities have been observed in most other cloned animals, which has been collectively termed ‘cloning syndrome’ ^23^. In total, *DAZL^-/-^* lambs displayed developmental malformations beyond the germ cell loss which resemble the cloning phenotype syndrome.

We have shown that testes from *DAZL^-/-^* male neonatal sheep are devoid of germ cells. *Dazl* is initially expressed around 11.5 days post-conception in inbreed C57BL/6 mice, where it is crucial for the correct acquisition of germ cell fate in both sexes ^13^. This specification role is likely conserved in sheep, since *DAZL^-/-^* neonatal male lambs lack germ cells. In female sheep, the germ cells begin to enter meiosis around 55 days post-conception ^33^. If ovine *DAZL* provides a permissive state for germ cells to sexually differentiate, similar to the role in murine germ cell development, its expression likely starts before day 55 in both sexes. Therefore, ovine *DAZL^-/-^* germ cells will have approximately 100 days of development before birth, compared to approximately 8 days for mice. This gives a considerably longer time in sheep for germ cell quality control by apoptosis, the mechanism known to destroy aberrant male germ cell clones ^34^ and *Dazl* mutant germ cells in inbreed C57BL/6 mice ^15^. Further studies would be required to confirm the temporal expression of *DAZL* in fetal sheep testes, as well as analysing *DAZL^-/-^* germ cells at the time of *DAZL* function, to confirm its conserved licensing role for germ cell specification.

In *Dazl^-/-^* mice, the timing of germ cell loss depends on their genetic background. In outbred mice, there were no significant differences in germ cell numbers compared to wild-type controls 9 days after birth ^34^, while in inbreed C57BL/6 mice less than a day old, only a few germ cells remained ^15^. In adult outbred *Dazl^-/-^* mice, most seminiferous tubules contained A-type spermatogonia but no spermatids, indicating that the surviving germ cells were unable to successfully complete meiosis. Therefore, the *DAZL*^-/-^ phenotype in male sheep more closely resembles C57BL/6 than outbreed mice, despite concerns that it may be difficult to draw robust conclusions on genotype-phenotype interactions from inbreed strains ^35^.

Selected somatic marker gene expression remained unchanged in *DAZL^-/-^* testes. Three key cell populations in the testes that enable spermatogenesis are the Sertoli cells, which directly support the germ cells, the Leydig cells, which are the interstitial steroidogenic cell lineage, and peritubular myoid cells, the outermost cells surrounding the seminiferous tubules ^35^. In mice, transcription factors *Gata4* ^36^ and *Nr5a1* ^37,38^ are key regulators of the Sertoli and Leydig cell lineages. *Hsd3b1* encodes an enzyme in the pathway for steroid production in Leydig cells ^39^. *Csf1* and *Gdnf* encode signalling proteins secreted by Leydig ^40^ and peritubular myoid, as well as Sertoli cells ^41,42^, respectively. Both growth factors are implicated in stimulating self-renewal of spermatogonial stem cells (SSC). Expression of these somatic markers was not affected in *DAZL^-/-^* lambs, indicative of an intact sperm cell niche. Our observations are consistent with data from *NANOS2*^-/-^ pigs, which showed normal testosterone production and sexual development in germ cell depleted males ^8^.

Specific germ cell transcript signatures were reduced or abolished in the two *DAZL^-/-^* lambs. We selected germ cell markers based on gene expression in neonate mice, which identified the target gene *DAZL*, along with *NANOS2* and *DDX4*, as germ cell markers ^43,44^. This was confirmed by *in situ* hybridisation colocalising *DAZL*, *NANOS2* and *DDX4* mRNA to germ cells in mice ^10,45,46^. *Dazl*-deficient germ cells retain gene expression characteristic of migrating PGCs, suggesting that *DAZL* is involved in committing pluripotent PGCs to the germ cell fate ^13,14,17^. Thus, we also surveyed expression of pluripotency-related genes *LIN28*, *SALL4* and *NANOG* in *DAZL*^-/-^ germ cells.

All germ cell markers were reduced by 1-2 orders of magnitude or below the detection limit relative to the wild type, concurrent with the morphological disappearance of spermatogonia from the testis cords. Since we did not observe any germ cells by histological and immunofluorescent analysis, we cannot attribute the remaining low-level *DAZL, NANOS2*, *DDX4,* and *SALL4* transcripts to any germ cells in *DAZL*^-/-^ testes. These data indicate that the somatic cells of the testis may produce a low-level background germ cell transcript signature or that there may be some rare germ cells remaining at the neonatal stage. The notion that tumorigenic germ cells may remain in *DAZL*^−/−^ gonads is supported by the observation that *DAZL*^−/−^ female pigs frequently form teratomas, which are absent from *DAZL*^+/+^ wild-type controls ^17^. The rate of teratoma formation was also significantly elevated in *DAZL*^−/−^ 129S mice ^17^, which naturally show incidences of spontaneous testicular teratomas between 1-3% ^47^. Likewise, DAZL mutations in humans were associated with an increased testicular teratoma risk ^48^. In contrast to pigs, humans and 129 mice, no teratomas were observed in several other *Dazl*^−/−^ mouse strains ^17^. Further research is required to determine if uncommitted PGCs are completely or incompletely eliminated by apoptosis in other *DAZL*^−/−^ mammals, including sheep, resulting in resistance or susceptibility, respectively, to teratoma formation ^49^. Testicular teratomas would limit the use of *DAZL*^−/−^ sires for agricultural breeding purposes due to animal welfare concerns.

The ovine *DAZL*^-/-^ germ cell-deficient phenotype confirms the important role of this gene in fetal germ cell development. In primates, autosomal *DAZL* transposed onto the Y-chromosome to produce several copies of a homolog, *DAZ* ^50^, which contains additional repeats of the DAZ domain, and is functionally redundant with murine *Dazl* ^51^. Deletions in the Y-chromosome region containing *DAZ* are associated with infertility in humans, accounting for around 13% of men with azoospermia ^52^. *DAZL* may also be implicated in female human infertility ^53^, opening up opportunities for genome-edited sheep models to study DAZL-related human pathobiology.

The overall results prove the hypothesis that the germline can be specifically ablated and therefore provide a niche for the development of germ cells from an elite donor. Refined understanding on the role of *DAZL* in germ cell development suggests that the optimal gene for producing germ cell-deficient sires should be further down the germ cell differentiation pathway. After *DAZL*-induced germ cell determination from uncommitted PGCs, teratoma formation from germ cell ablation would potentially be limited. This criterion identifies *NANOS2*, which plays a role post-germ cell determination, as a good target ^54^. For agricultural applications, germ cell-deficient sheep would provide a suitable host for absolute transmission of a donor germline in chimaeras, either from transplanted SSCs or from embryonic blastomeres developing into germ cells. The former approach relies on culturing molecularly and functionally well-defined bona fide sheep SSCs to repopulate the vacant seminiferous tubules and reconstitute normal sperm output ^55^. This is a considerable limitation that has so far not been resolved in livestock. The latter approach would generate chimaeras by embryo complementation, a mechanism that has already been demonstrated as feasible in sheep ^56^, cattle ^9,57–59^ and pigs ^60–62^. *NANOS3*^-/-^ germline-deficient female cattle embryos have been used as hosts in chimaeras ^9^, but further work would be required to unequivocally attribute embryo-derived germ cells to the donor germline. This approach would also need to be progressed in males, which are more relevant for livestock breeding since they generate far more offspring, either by natural mating or artificial insemination. Ultimately, the semen of *DAZL*^-/-^ rams at full adulthood needs to be assessed confirm the complete absence of spermatozoa to ensure there would be no contaminating endogenous spermatogenesis in a chimaera. The generation of ‘absolute transmitters’ from complementing genetically sterilised livestock hosts with genomically-selected elite embryonic donor cells provides an exciting opportunity for accelerating genetic gain ^63^.

## Materials and methods

Investigations complied with the New Zealand Animal Welfare Act 1999 and were approved by the Ruakura Animal Ethics Committee.

### Donor cells

Ovine fetal fibroblasts (OFFs) were established from a male slaughterhouse fetus (crown-rump length: 230 mm, estimated age 12-14 weeks of gestation) following our standard operating procedures in cattle ^64^. Briefly, the skin was dissected, washed briefly in 70% ethanol, minced, and cultured in hanging drops. Genotyping using an in-house ovine high-density SNP chip indicated that the resulting line (‘OFF3’) was a composite of Perendale, Texel, Coopworth and Romney breeds, in descending order of relative contribution (K. Dodds, personal communication). For somatic cell cloning, G_0_ cells were obtained by culture in medium containing 0.5% FCS for 5 days and harvested by trypsinization ^64^.

### Genome editing

The sequence of the *DAZL* genes obtained from NCBI databases from sheep, mouse and cattle were compared. The predicted mRNA (sheep: XM_027964130.1, mouse: NM_010021.5, cattle: NM_001081725.1) and coding sequences (sheep: XP_027819931.1, mouse: NP_034151.3, cattle: NP_001075194.1) were aligned.

Several gRNAs were identified within the target region for *DAZL* and selected for minimal off-target sites using CCTop (Supplementary Table S1). Expression vector pSpCas9(BB)-2A-Puro (PX459) V2.0, a gift from Feng Zhang (Addgene plasmid # 62988; http://n2t.net/addgene:62988; RRID:Addgene_62988), was used for transcription of gRNA, Cas9 and puromycin resistance ^65^. A single-stranded oligonucleotide with 50 bp arms for HDR (Supplementary Table S1) was also delivered with PX459 using the Neon® Transfection System (Thermo Fisher, New Zealand) at 1,500 V with one 20 ms pulse. After selection with puromycin (2 µg/mL) for 48 h, genomic DNA was isolated. The QX200 (Bio-Rad) ddPCR system was used to quantify HDR using hydrolysis probes for both the wild-type and mutant variants with VIC and 6-FAM fluorophores, respectively (Supplementary Table S1).

### Manual isolation of clonal cell strains

The population with the highest proportion of HDR events was used for manual isolation of clonal cell strains. We modified our mitotic shake-off procedure for synchronising cells in the G1 cell cycle stage to obtain sufficient starting material for isolating genome-edited clones ^66,67^. Briefly, 2.5×10^4^ cells/cm^2^ were seeded on 4-well plate and cultured for 18-20 h. Cells were washed once with PBS and cultured for another 2-3 h before mitotic shake-off by gently tapping the dish with moderate horizontal force. Individual mitotic cells or couplets in ana- or telophase were selected with a finely pulled glass Pasteur pipette and seeded into single wells of a 96-well plate. Cells were expanded for 6-7 days until a confluent colony formed, passaged onto a 48 well plate, and cryopreserved once confluency was reached.

### Validation of genome editing

Genomic DNA was isolated from cell lines in lysis buffer (100 mM Tris pH 8, 1 mM EDTA, 0.5% (v/v) Tween-20, 0.5% (v/v) Triton X-100) and 1 mg/ml Proteinase K (QIAGEN, Germany). The solution was incubated at 55°C for 15 min, and heat inactivated by incubating for 5 min at 95 °C. We used the Genotyping ToughMix (Quantabio, USA), 0.9 µM primers, 0.25 µM of the hydrolysis probes and 1 µl of template up to a final volume of 10 µl. For RT-qPCR, samples were run on the Rotor-Gene 6000 (Corbett Life Science, Australia) with the following program: initial denaturation at 95°C for 5 min; then 40 cycles of 95°C for 15 seconds, 60°C for 60 seconds. For the resulting Cq value, copies were calculated from a standard curve that was prepared by diluting genomic DNA of known concentration for the wild-type allele, and a synthetic gBlock (Integrated DNA Technologies, USA) template for the mutant allele.

For identified mutant clones, a larger genomic region was PCR-amplified (Table S1) and digested with 1 unit of *Taq*I at 65°C for 4 h. Insertion was confirmed by Sanger sequencing (Massey University Genome Service, New Zealand) of PCR-amplified fragments, using above primers (Supplementary Table S1).

### Somatic cell cloning, pregnancy monitoring and parturition

Sheep somatic cell cloning was carried out based on a modified protocol for cattle zona-free embryo reconstruction ^64^. Briefly, zona-free MII oocytes were enucleated approximately 19 h after *in vitro* maturation, and serum-starved OFF3 donor cells were attached in 10 µg/mL phytohemagglutinin P (PHA-P), then fused by automatic alignment using ∼80 V/cm alternating current before two direct current pulses of 2 kV/cm for 10 µsec in 270 mOsm fusion buffer (260 mM D-mannitol, 0.05 mM CaCl_2_, 0.1 mM MgSO_4_, 0.5 mM Hepes, 0.05% fatty acid-free bovine serum albumin). A second cytoplast was attached in PHA-P and fused at 1.5 kV/cm to produce a double-cytoplast reconstruct ^68^. Activation by ionomycin (5 μM; 4.5 min)/6-dimethylaminopurine (2 mM; 4 h) was carried out 3-4 h post-fusion. Groups of 10 embryos were cultured in 20 µl drops of SOFaaBSA, each embryo occupying a manually prepared micro-well ^69^. After five days, embryos were transferred into single culture with individual 5 µl SOFaaBSA drops. Blastocysts were morphologically graded on day 7 according to the International Embryo Technology Society guidelines ^70^ and either vitrified or transferred into recipient ewes by laparoscopic surgery. Vitrification and warming was performed as previously described ^71^. Pregnancies were monitored by ultrasound throughout gestation and hormonal induction carried out with 16 mg dexamethasone administered twice on day 145 for delivery on day 147.

### Testes histology

Neonate testes from edited and wild-type animals were collected and either fixed in Davidson’s fixative for paraffin embedding, fixed in 4% paraformaldehyde (PFA) with 4% (w/v) sucrose for frozen sections, or the testis parenchyma was snap-frozen in liquid nitrogen for gene expression analysis. Davidson’s-fixed testes were embedded in paraffin, sectioned, haematoxylin–eosin stained, analysed by a pathologist service (Veterinary Pathology Ltd., Hamilton, New Zealand) and photographed. Cryosections we made by washing PFA-fixed testes through 30% (w/v) sucrose, freezing in Optimal Cutting Temperature (OCT) compound, then sectioned at 8 µm using a Leica CM1850 cryostat (Leica Biosystems). The OCT sections were immuno-stained by permeabilising in 0.1% Triton X-100 for 10 min, quenched with 100 mM NH_4_Cl for 10 min, blocked in 5% BSA for 20 min, incubated in 1:100 anti-DDX4 (ab13840, Abcam) overnight at 4°C, and labelled with anti-rabbit secondary antibody Alexa Fluor® 568 (A-11036, Thermo Fisher Scientific) containing 5 µg/ml Hoechst 33258. Images were taken with an epifluorescence Olympus AX70 microscope and processed with ImageJ (version 1.45s; National Institutes of Health) for identical background subtraction and colourisation of wild-type and mutant samples.

### Gene expression analysis

Snap-frozen testis parenchyma was homogenised in TRIzol Reagent (Thermo Fisher Scientific) and RNA extracted following the manufacturer’s instructions. Potentially contaminating genomic DNA was removed with DNase I kit (Thermo Fisher Scientific), with an incubation for 30 min at 37°C. A sample was omitted for subsequent reverse transcription as a control to ensure no genomic DNA was present. cDNA was synthesised using SuperScript® IV (Thermo Fisher Scientific), processed following the manufacturer’s instructions. KAPA SYBR FAST qPCR Kit (Kapa Biosystems) was used with a MIC qPCR Cycler (Bio Molecular Systems), the 10 µl reaction mix consisted of KAPA SYBR FAST master mix, 0.8 µM of each primer and 1 µl of 1:4 diluted cDNA template. A ‘no template’ control was run in each experiment, and samples were run in triplicates. The following program was used: 1) denaturation (30 sec at 95°C); 2) amplification and quantification (5 sec at 95°C, 30 sec at 60°C, followed by 30 sec at 72°C with a single fluorescent measurement repeated 40 times); 3) melting curve (95°C, then cooling to 70°C for 1 min, heating at 0.1°C/sec to 95°C while measuring fluorescence), unless indicated otherwise (Supplementary Table S2).

For reference genes, we relied on a previous study that identified *GAPDH*, *HPRT* and *ACTB* as stably expressed in testes tissue (PMID: 24952483). For markers of spermatogonial and somatic cell populations, we relied on a single cell qPCR screen at the neonatal stage in the mouse ^44^. After designing primers for 17 genes using NCBI Primer-BLAST, spanning introns when possible, we validated their expression in wild-type testis by confirming i) amplification of a single product through gel electrophoresis, and ii) sufficient dynamic range over four orders of magnitude by running a 4-fold dilution series. Only the primer pairs that passed these quality controls were used for analysis, which resulted in *DAZL*, *NANOS2*, *DDX4*, *SALL4*, *NANOG*, and *LIN28* as spermatogonial markers and *GATA4*, *HSD3B1*, *NR5a1*, *CSF1,* and *GDNF* as somatic markers (Supplementary Table S2). Assays were optimised to ensure a single melting peak corresponded to the correct PCR product size and absence of primer-dimer formation.

Relative quantification was carried out as described ^72^, taking reaction efficiency into account by amplification curve analysis. The mean reaction efficiencies for each assay is listed in Supplementary Table S2. When amplicons were not detected (as indicated by a single specific melting peak) an arbitrary baseline was set for that sample with a Cq of 35, which was then normalised on reference genes.

### Statistical Analysis

An Analysis of Variance Model (ANOVA) was used for analysing ddPCR quantification of genome editing efficiency ^73^. For qPCR gene expression analysis, log transformed relative expression was analysed by linear mixed-effects model fit by REML ^74^. A Cq value of 35 was given for missing values for computation of the baseline relative expression.

## Acknowledgements

We thank Brigid Brophy for Cas9 primers, Pavla Turner for assistance with cloning, Alan Julian for histological services, Stephanie Delaney for animal husbandry, Alison Cullum for veterinarian care, and Harold Henderson for assistance with statistics. ZM was supported by Callaghan Innovation R&D student fellowship grant, AgResearch, Livestock Improvement Corporation fellowship grant, Todd Foundation Award for Excellence, and Heritage Incorporated Magnus Grant. SJA was supported by University of Auckland Doctoral Scholarship and Todd Foundation Award for Excellence. This research was supported by funds from AgResearch and University of Auckland.

## Author contributions

Conceived and designed the experiments: ZM, BO, RGS. Performed the experiments: ZM, SJA, JW. Analyzed the data: ZM, BO, RGS. Contributed reagents/materials/analysis tools: ZM, SJA. Wrote the paper: ZM, BO, SJA, RGS.

## Additional Information

### Competing financial interest

JW and BO are employees of AgResearch. There are no patents, products in development or marketed products to declare.

## Supplementary Figures

**Supplementary figure 1.**
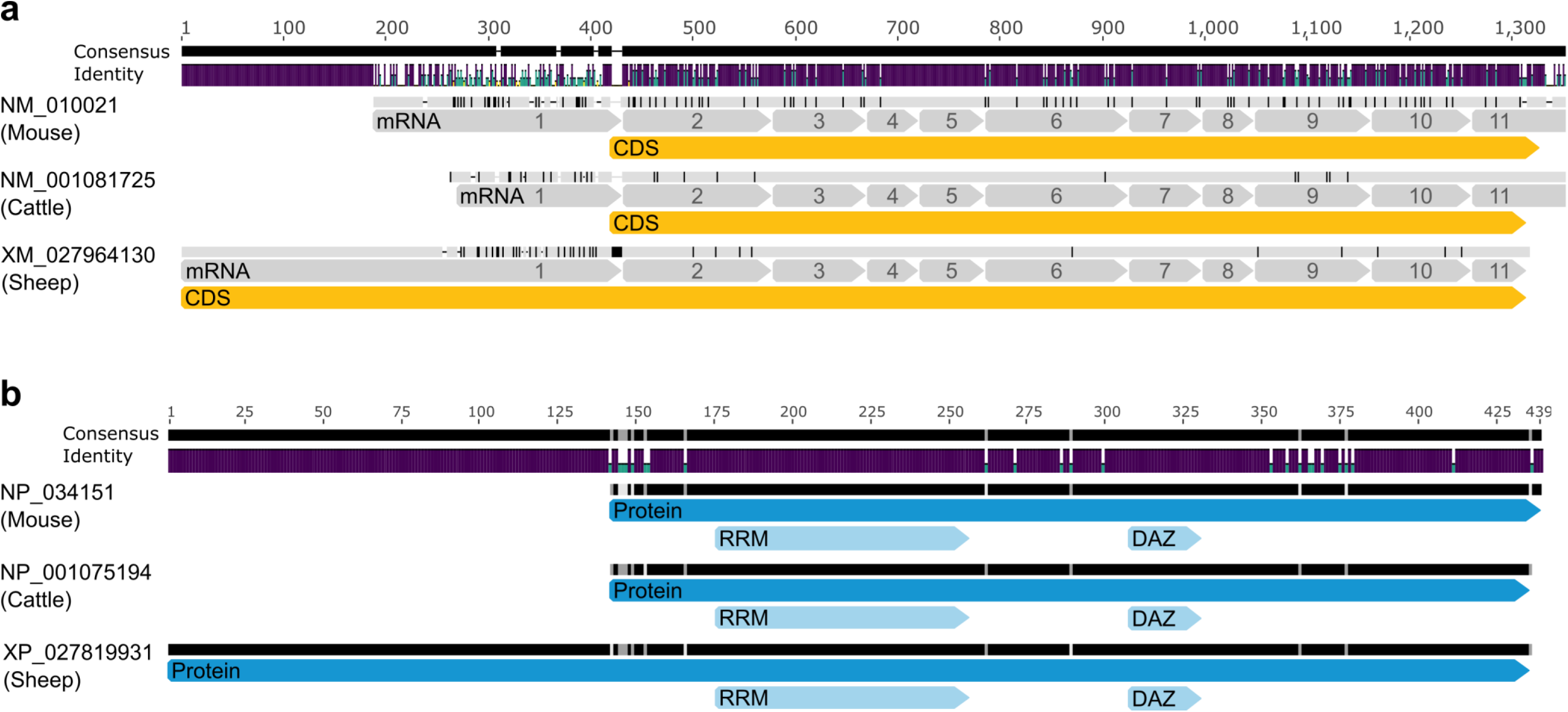
Sheep, mouse and cattle mRNA and protein alignments for *DAZL*. (a) Alignment of mRNA sequences showing the consensus areas with insertions indicated by dashes, and the proportion of identity between sequences shown by colour (with purple for complete, green for moderate, and yellow for little identity). Species-specific sequence differences are shown with horizontal and vertical bars, denoting insertions and different identity, respectively. The mRNA sequence (grey) is broken up into numbered exons and the coding sequence (CDS) is indicated in yellow. (b) The visualisation of the protein alignment has the same variables as the mRNA alignment, with the length of the amino acid sequence for each species indicated in blue and the functional RNA recognition motif (RRM) and DAZ domain given in light blue.

**Supplementary figure 2.**
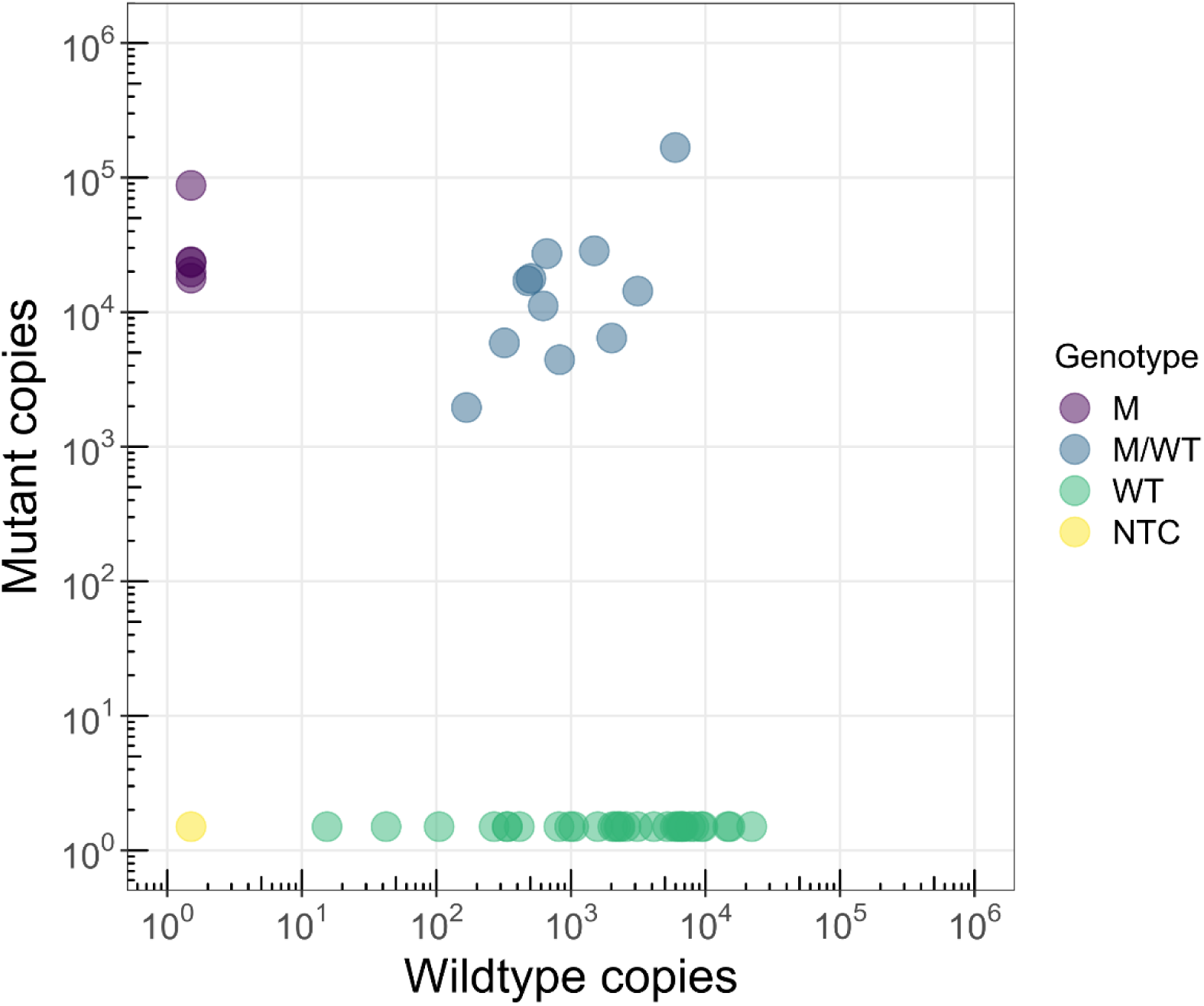
Allelic discrimination assay for identification of biallelically edited strains. The separation of strains into mutant (M), heterozygous (M/WT), wildtype (WT), and no template control (NTC) is according to the presence of the mutant or wild-type allele determined by qPCR.

**Supplementary figure 3.**
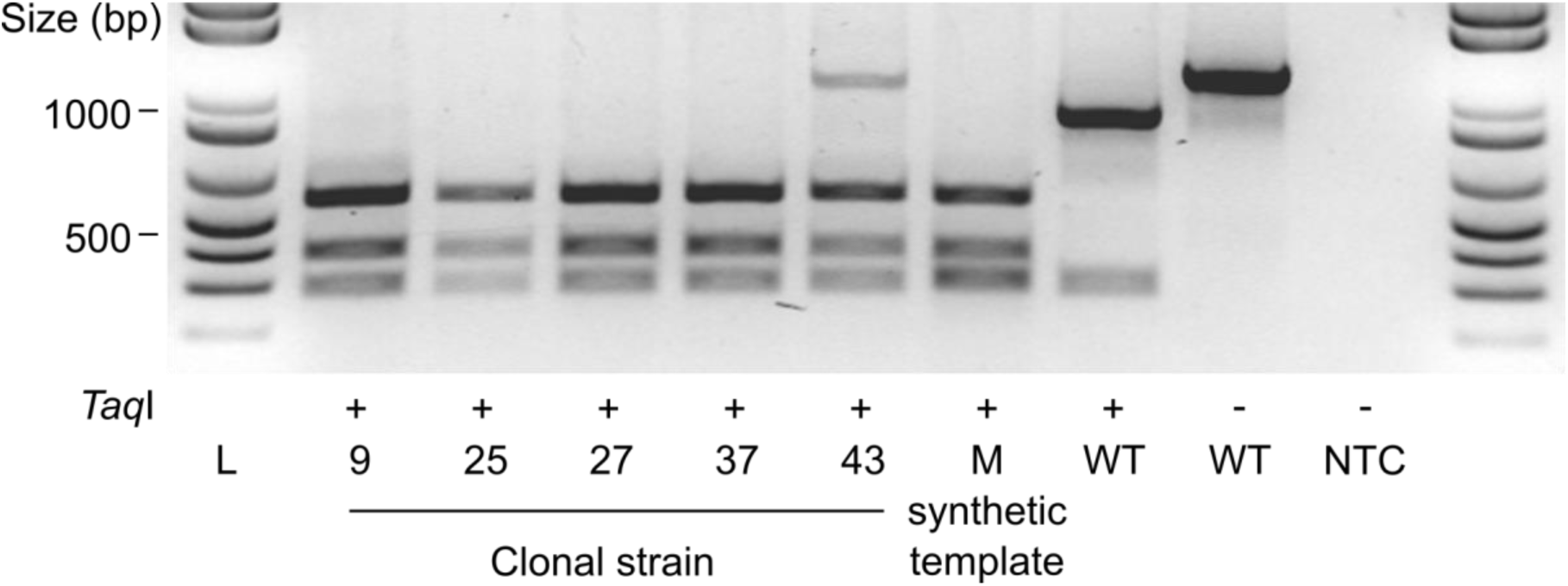
*Taq*I digest for validation of HDR mediated insertion. PCR amplicons of strains were exposed to *Taq*I, along with a synthetic mutant (M), wildtype (WT) and no template control (NTC). Presence or absence of *Taq*I is indicated by + or -, respectively.

**Supplementary figure 4.**
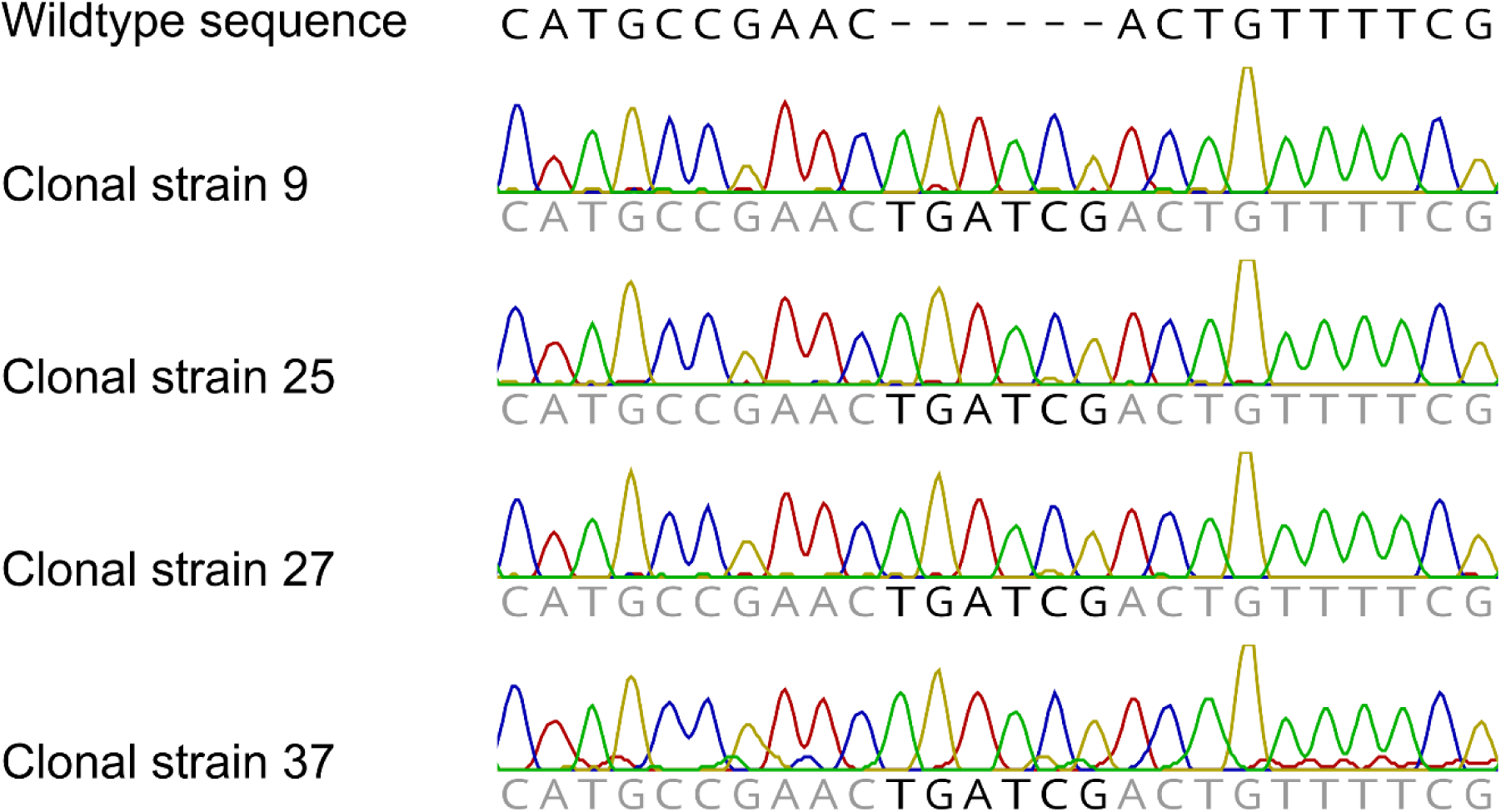
Sanger sequencing chromatograms confirming editing of strains. HDR-mediated insertion is highlighted by black text

**Supplementary figure 5.**
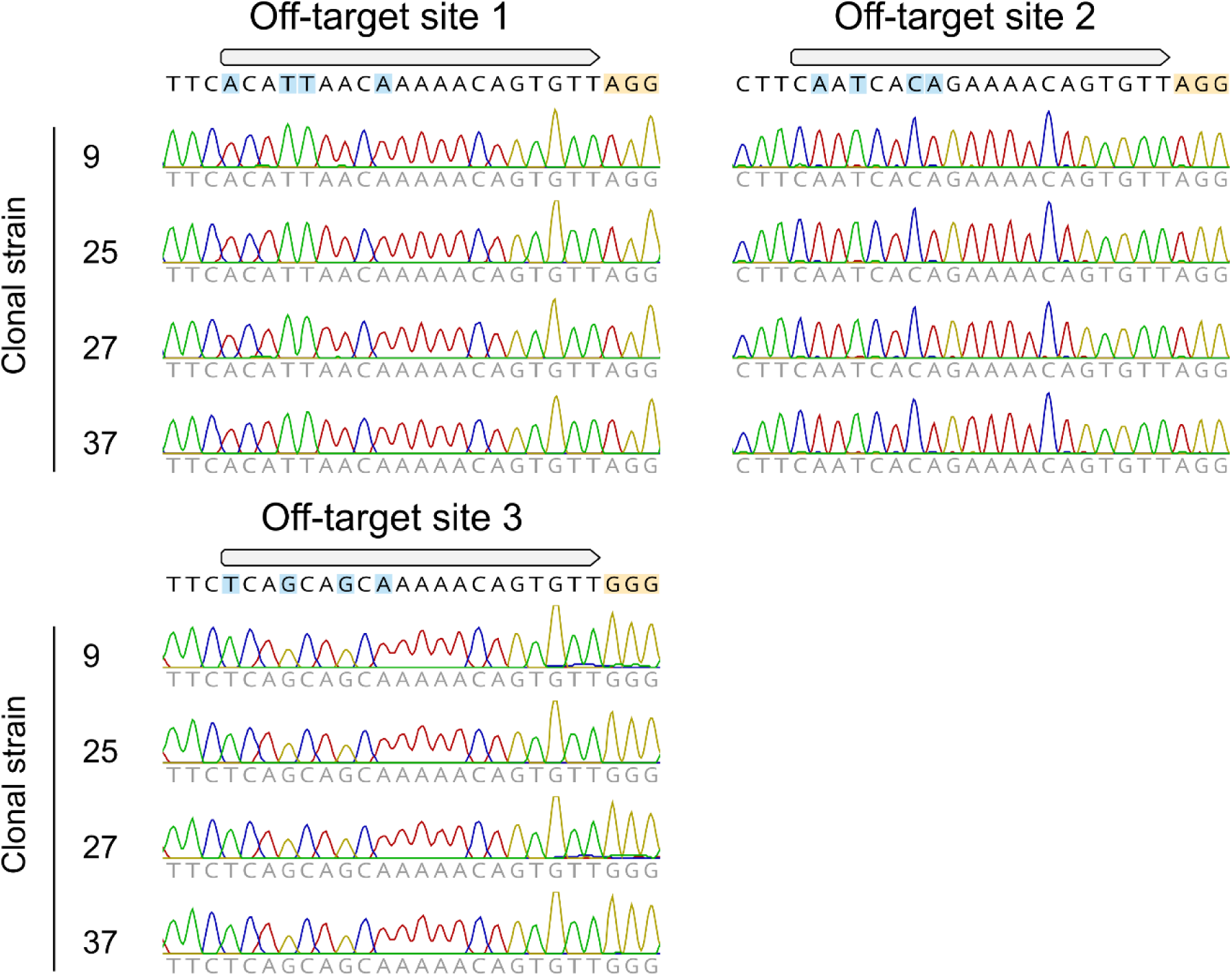
No mutations revealed in Cas9 off-target analysis. Sanger sequencing each strain for the top three off-target sites identified by CCTop. The nucleotide mismatches to the gRNA are highlighted in blue, and the PAM sequence in orange.

**Supplementary figure 6.**
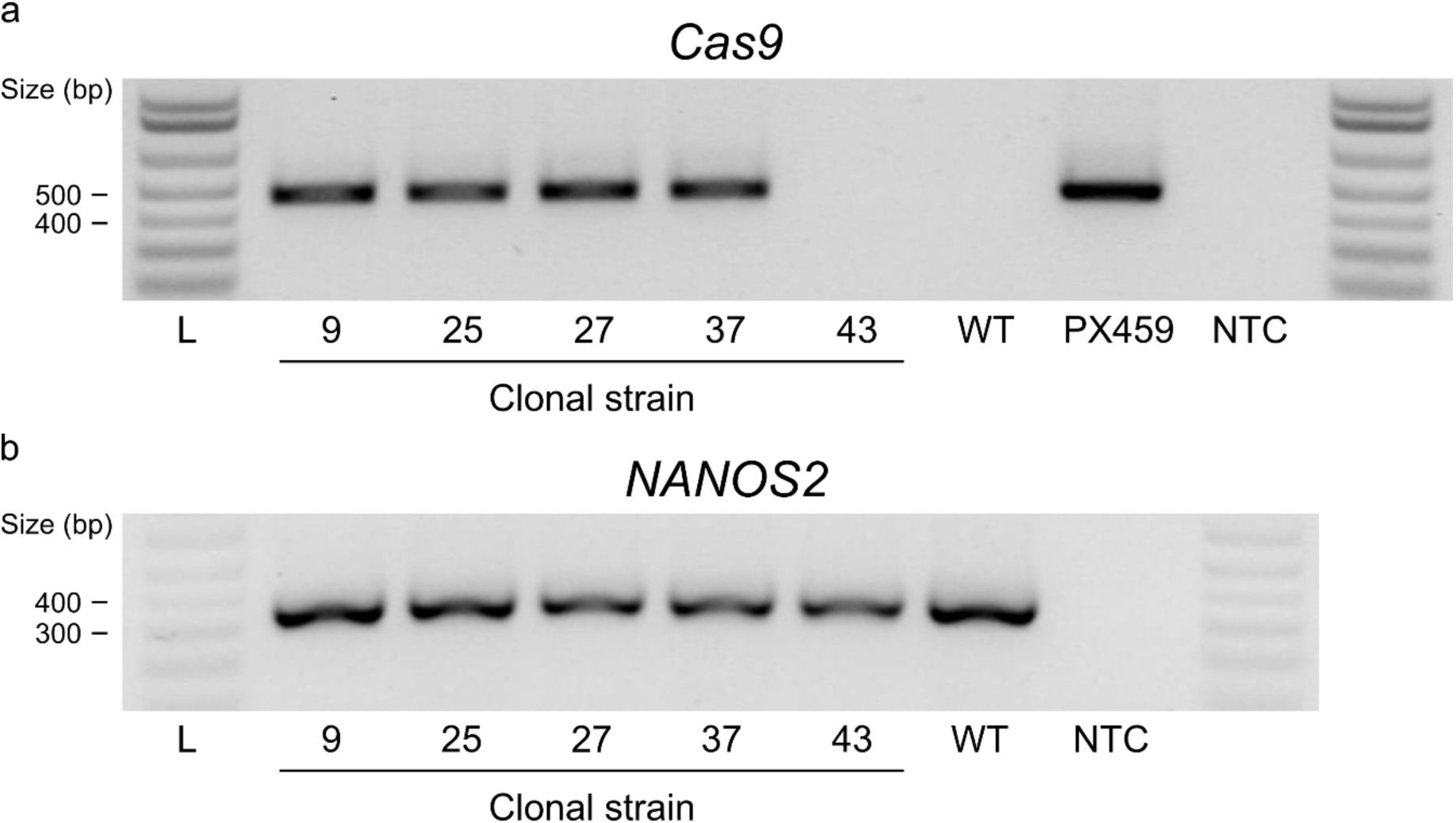
gRNA-Cas9 plasmid integration in strains. (a) PCR amplification of *Cas9* coding sequence in strains, wild-type parental cells (WT), the PX459 gRNA-Cas9 plasmid, and a no template control (NTC). (b) Presence of genomic DNA was validated by amplification of genomic *NANOS2* region.

**Supplementary figure 7.**
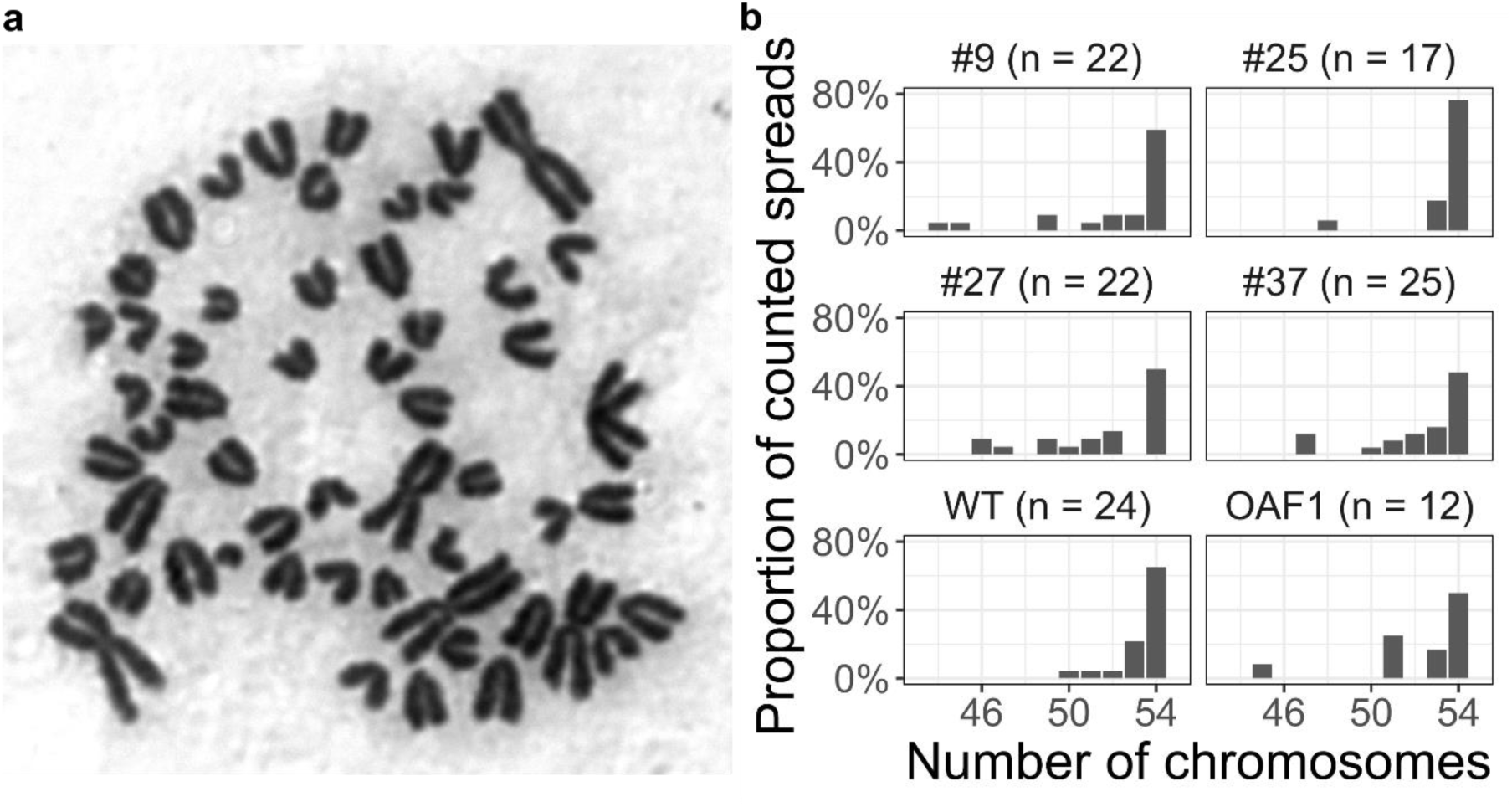
Karyotype confirmed in strains. (a) Representative spread with 54 chromosomes. (b) Chromosomes counted in spreads of strains. Chromosomes were also counted in early passage wild-type OFF3 parental cells (WT) and OAF1 cells which have produced lambs after somatic cell cloning.

**Supplementary figure 8.**
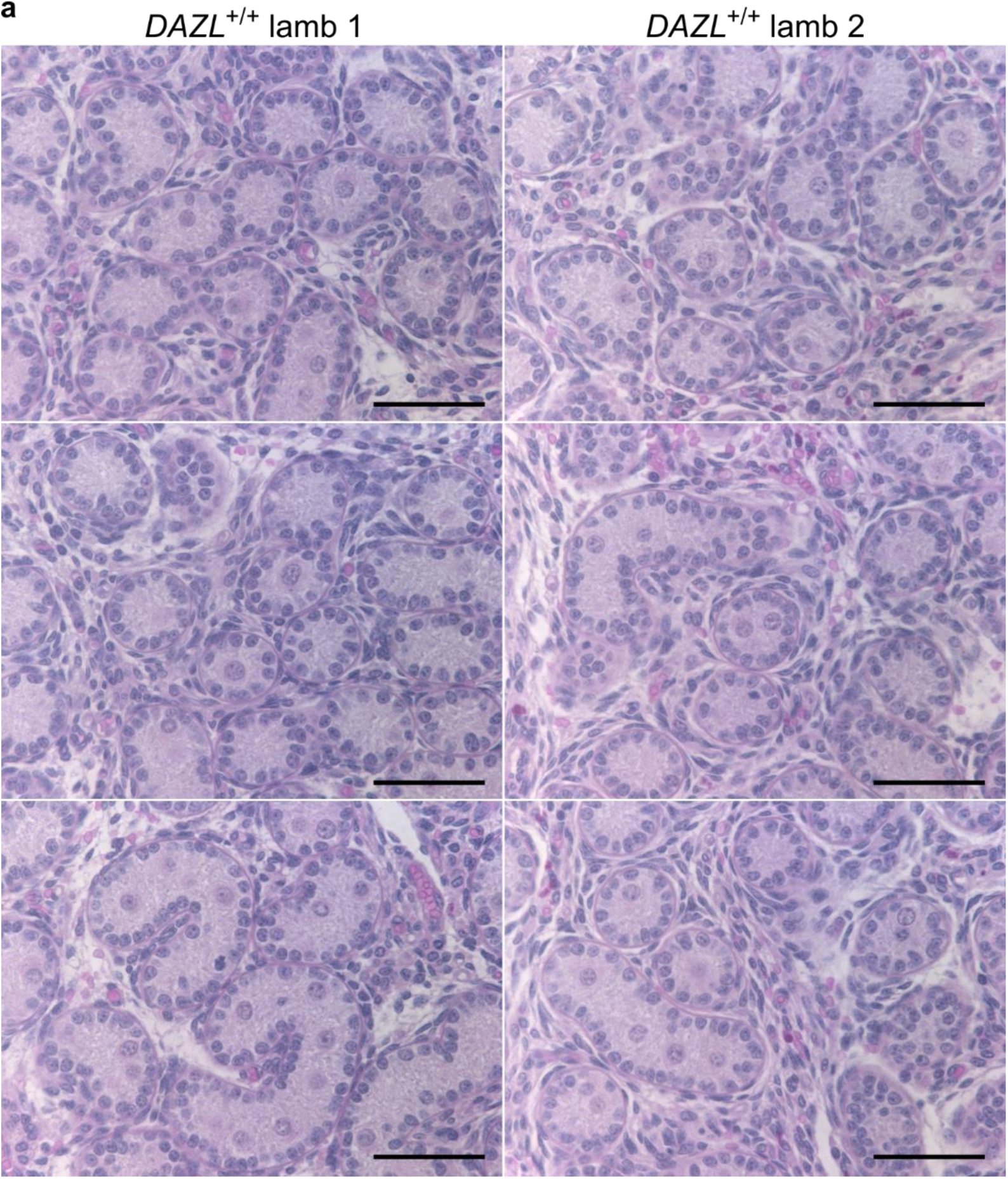
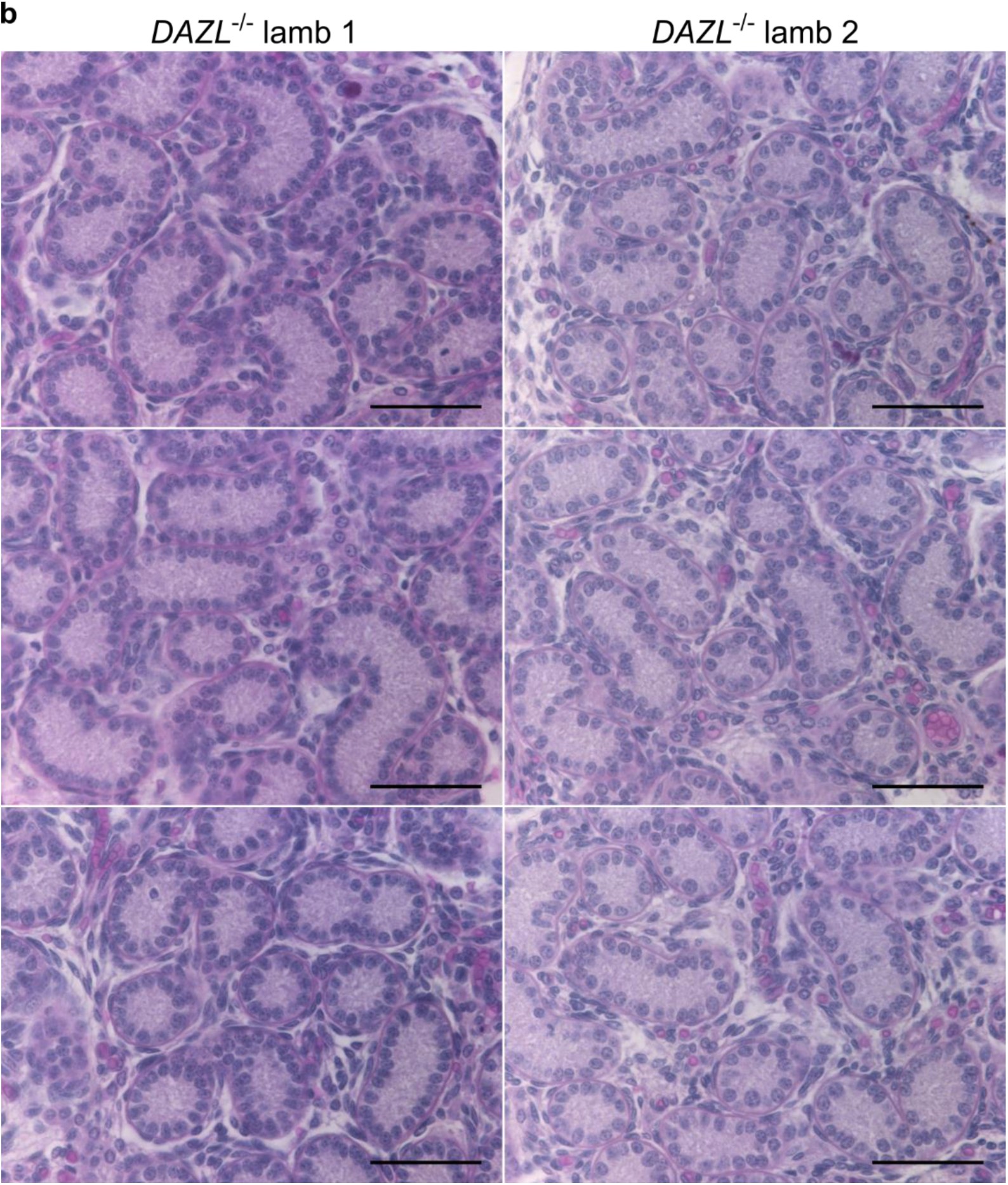
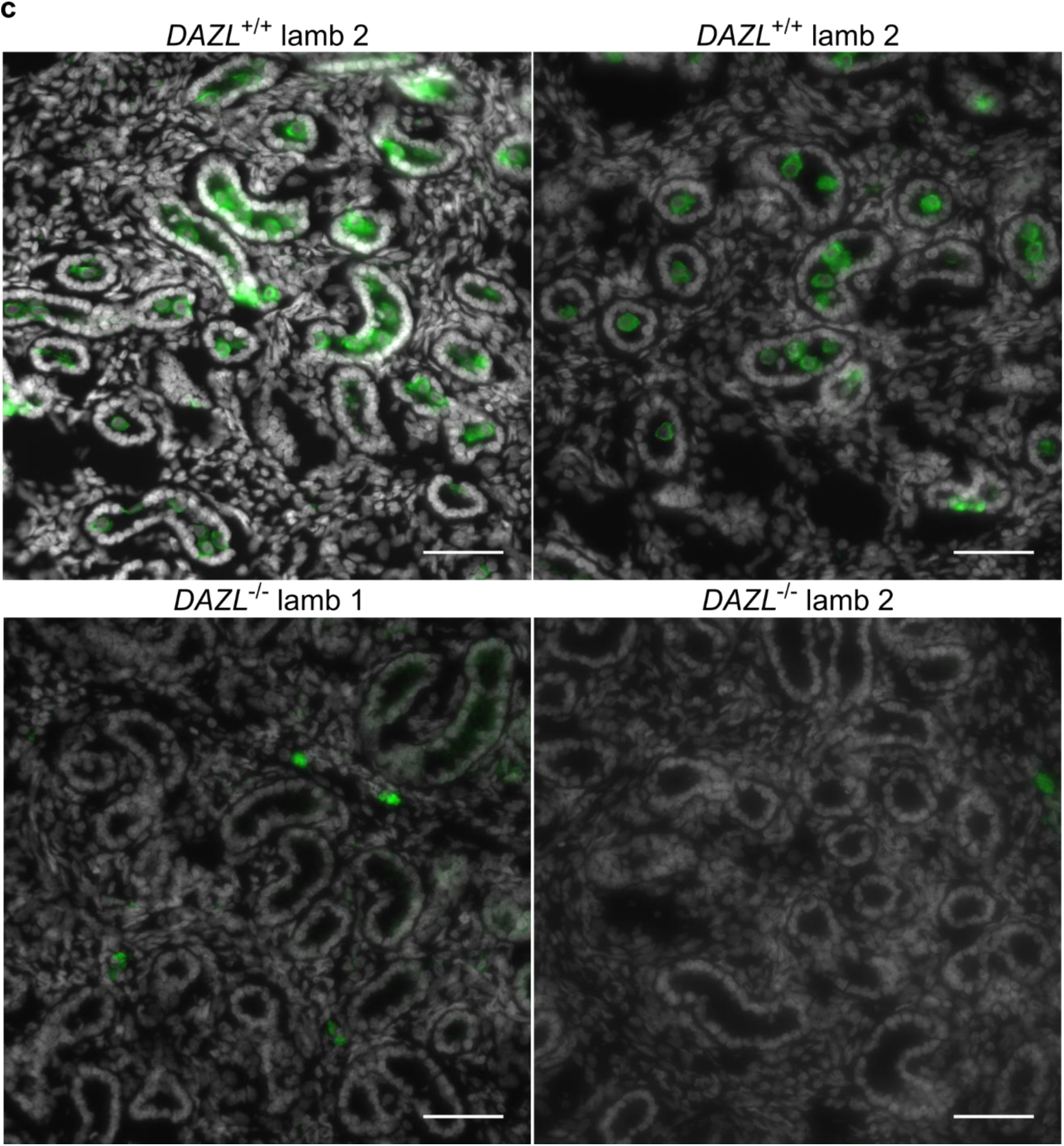
Additional images of germ cell-deficient phenotype in *DAZL*^-/-^ neonate testis. Haematoxylin and eosin stained sections of (a) *DAZL*^+/+^ and (b) *DAZL*^-/-^ neonate lamb testis. (c) Immunostaining of DDX4 (green) in *DAZL*^+/+^ and *DAZL*^-/-^ neonate lamb cryosections, co-stained for DNA (Hoechst 33342). Scale bars are 50 µm.

## Supplementary tables

**Supplementary table 1.**
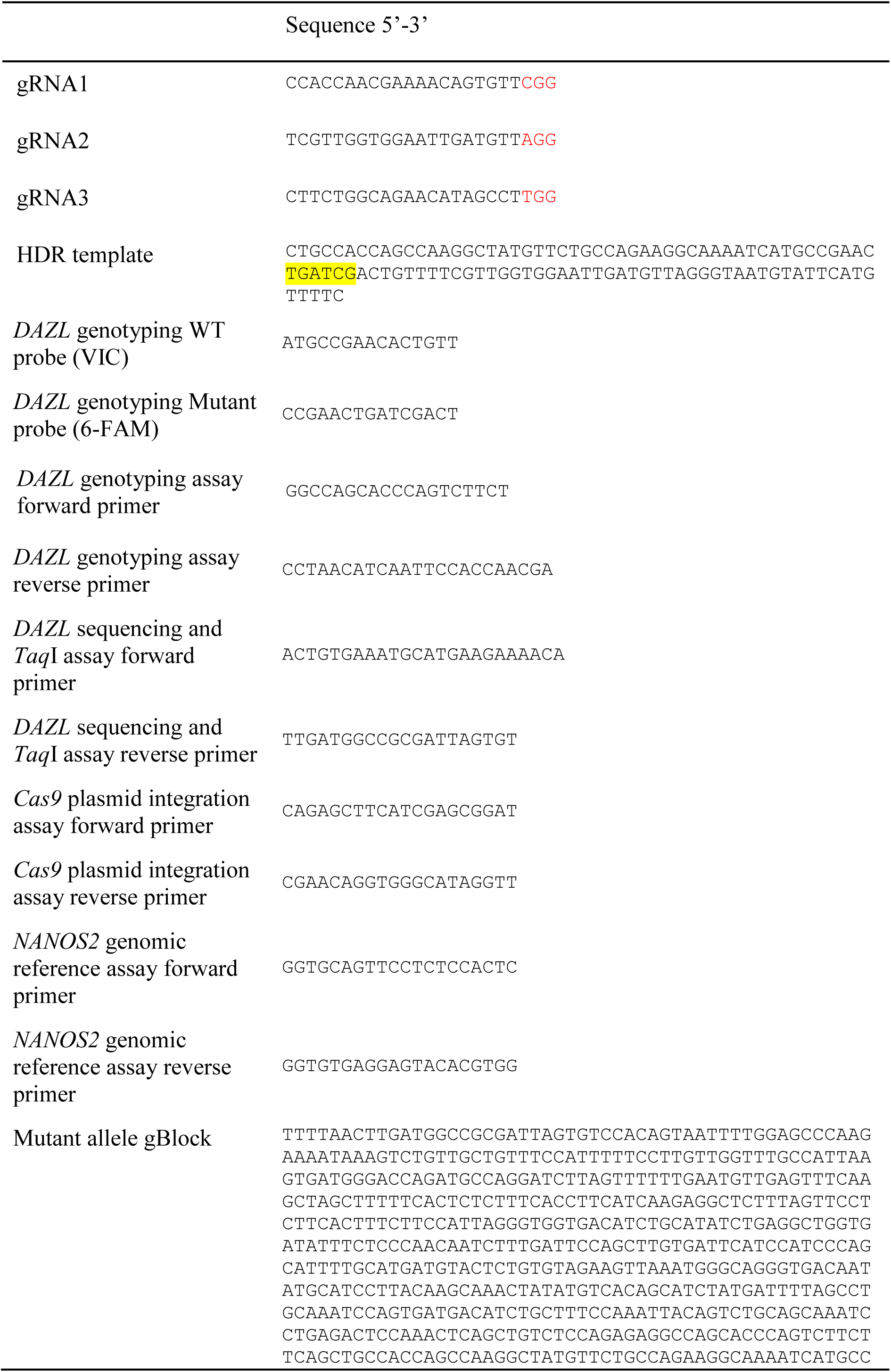
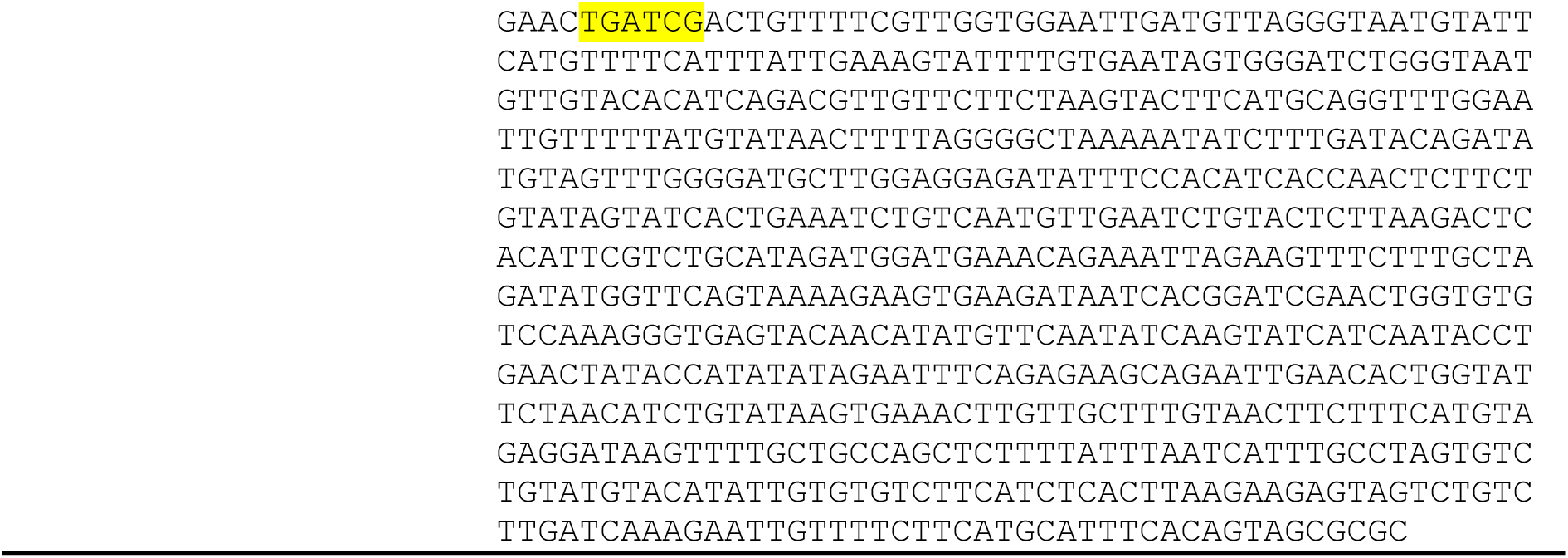
Sequences for genome editing tools. The PAM site of the gRNA is highlighted in red text. The 6 bp insertion within the HDR oligonucleotide and mutant allele gBlock are highlighted in yellow.

**Supplementary table 2.**
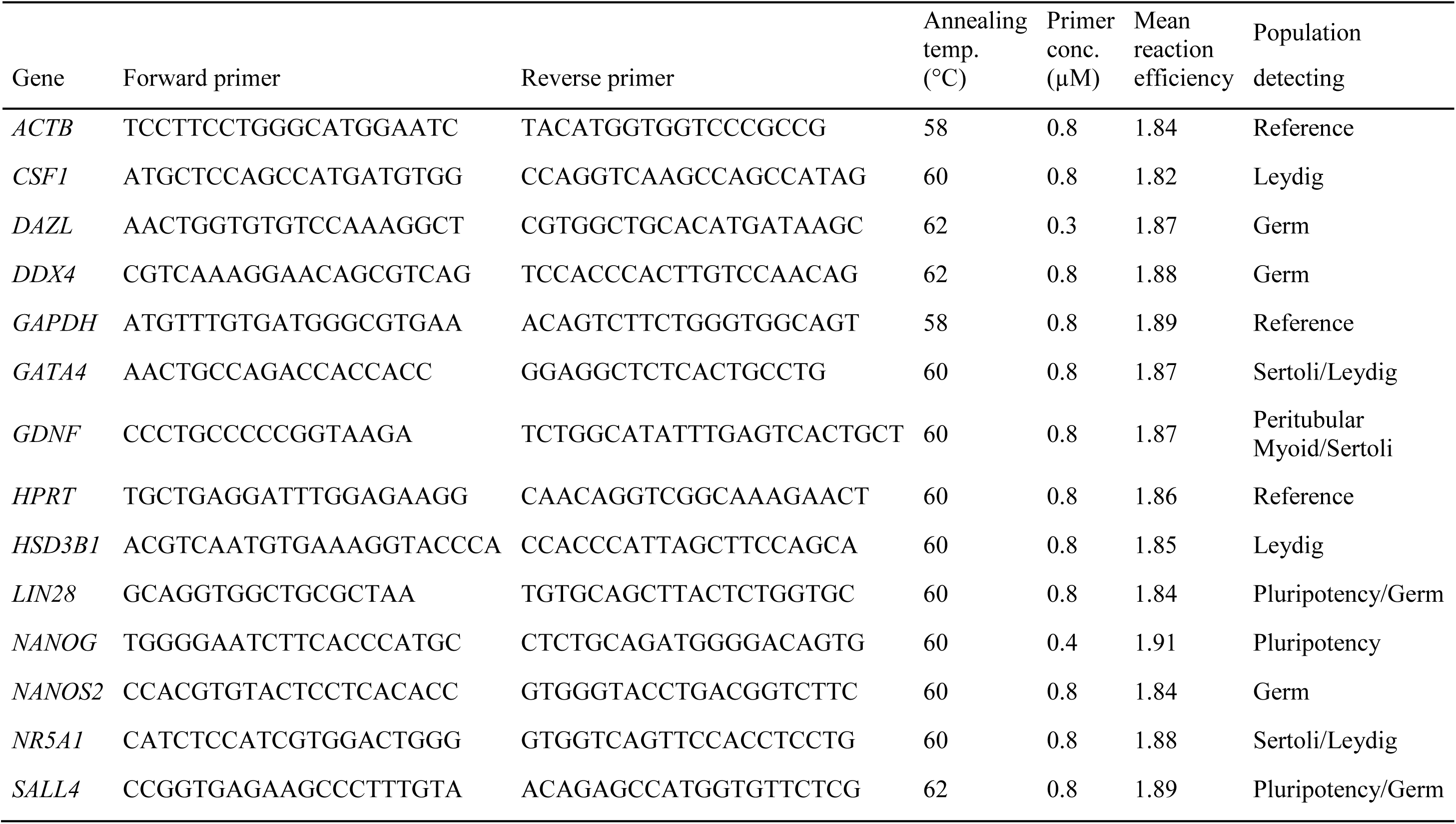
Gene expression primer sequences and reaction conditions.

## Notes

#### Summary of Updates

Introduction and discussion revised to engage with Nicholls et al. (2019), published after the submission of our initial manuscript.

